# The response of the relationship between carbon components and soil aggregates in vegetable fields to continuous input of different carbon sources

**DOI:** 10.1101/2025.04.02.646932

**Authors:** Yuan Yang Peng, Jiang Ling Huang, Yan Qin Peng, Yin Peng, Shi Xiong Li, Xi la tu Da Bu

## Abstract

**Aims:** To address the issue of declining soil organic carbon (SOC) in vegetable fields, this study investigated the effects of continuous input of different carbon sources on soil carbon pool dynamics. Methods:A five-year field plot experiment was conducted to analyze the impact of organic fertilizer, biochar, and their combined application on soil organic carbon components, aggregate stability, and carbon pool characteristics. Results:The continuous application of biochar and organic fertilizer significantly increased the content of soil organic carbon components. Organic fertilizer contributed more actively to labile organic carbon fractions, whereas biochar primarily supplied stable organic carbon. All treatments enhanced the proportion of macroaggregates (>0.25 mm) and improved aggregate stability, with organic fertilizer exhibiting superior effects compared to biochar. Organic fertilizer also promoted organic carbon sequestration in aggregates by enriching readily oxidizable organic carbon (ROC) and water-soluble organic carbon (DOC). In contrast, biochar application significantly increased inert organic carbon (IOC), leading to higher total SOC compared to organic fertilizer alone. Carbon pool characteristic indices improved significantly under continuous biochar and organic fertilizer application. Organic fertilizer treatment showed a higher carbon pool activity index, while biochar outperformed in other indices. The combined application yielded the most favorable results. Correlation analysis revealed that soil aggregate stability was significantly associated with DOC and microbial biomass carbon (MBC). Both biochar and organic fertilizer enhanced soil available nutrient content, which increased dynamically over time. Principal component analysis confirmed that the combined application of organic fertilizer and biochar was most effective for soil carbon sequestration and fertility improvement. Conclusion:Biochar and organic fertilizer differentially influenced soil organic carbon composition. Organic fertilizer improved carbon pool characteristics by mediating macroaggregate formation through labile organic carbon (e.g., DOC and MBC), while biochar contributed directly to carbon sequestration due to its inert and stable nature. Their combined application synergistically enhanced the formation and stability of soil macroaggregates, improved soil carbon pool characteristics, and optimized nutrient availability in vegetable fields. This version improves clarity, conciseness, and flow while maintaining scientific accuracy. Let me know if you’d like any further refinements!

**Highlights:** The continuous application of biochar and organic fertilizer changed soil carbon components, enhanced aggregate stability and soil carbon pool characteristics.

Biochar and organic fertilizer increased soil available nutrient content, enhanced soil carbon sequestration and fertility.

The continuous application of biochar and organic fertilizer changed the carbon components in the aggregates, and the influence mechanism was different. The combined application had synergistic effect.

## 1 Introduction

Soil organic carbon composition, aggregates and their stability are important indicators to measure soil fertility. Aggregate is the basic unit of soil structure and composition, and it is also an important place for the existence of organic carbon in soil. Its quantity and quality determine many physical and chemical properties and fertility status of soil (Liu et al., 2011). Soil organic carbon and aggregates affect each other and are closely related. Organic carbon can not only enhance the agglomeration between soil particles, but also promote the formation of aggregates. Aggregates give physical protection to organic carbon, which can affect the characteristics of soil carbon pool(Dhaliwal et al., 2020).

Long-term single application of chemical fertilizers will destroy the structure of soil aggregates and accelerate the decomposition of organic carbon in aggregates, which is not conducive to carbon sequestration (Song et al., 2024). Exogenous addition of biochar and organic fertilizer is an effective measure to supplement soil carbon and improve the stability of soil aggregates (Lin et al., 2019). The application of organic fertilizer significantly increased the proportion of macroaggregates in soil and the content of dissolved organic carbon (DOC) in this aggregate, as well as the content of carbon, nitrogen and phosphorus in each particle size aggregate, thereby promoting soil function and crop productivity (Tian et al., 2022; Sun et al., 2023).(Chen et al., 2022) found that single application of biochar could significantly increase soil total organic carbon (SOC) content. The combined application of AM fungi and biochar had no significant effect on medium soil aggregates, but it could significantly increase DOC content and significantly reduce readily oxidizable organic carbon (ROC) content. (Li et al., 2017) found that single application of biochar or combined application of biochar and organic fertilizer could significantly increase the content of POC and ROC in soil, and also contribute to the growth and yield of apple plants. Under the combined application of organic fertilizer and biochar, the degree of soil humification was higher when biochar was added alone, and microbial biomass organic carbon (MBC) and DOC had a significant effect on the cumulative emission of soil CO2 (Zheng et al., 2019).(Zhang et al., 2023) found that single application of biochar could significantly increase the content of carbon components in soil and the content of carbon components in aggregates of different particle sizes through the positioning test of paddy soil. There is a significant correlation between the stability of soil aggregates and SOC storage (Hou et al., 2015). These results revealed that the distribution of SOC and its fractions in soil aggregates was affected by many factors, In addition, the active organic carbon in the soil carbon pool has strong environmental sensitivity and plays a key role in the transformation of soil physical, chemical and biological characteristics. It further highlights the necessity of in-depth study of the relationship between soil aggregates and SOC, in order to better understand the mechanism of soil SOC accumulation and transformation, and guide vegetable field management and soil protection measures. After the application of organic fertilizer, the efficiency of soil carbon sequestration is not high, the accumulation of organic carbon is not clear, and the mechanism of microbial turnover on the formation of aggregates is not clear. Biochar is mainly due to its stable physical properties and high stability after being applied to the soil, and the combination of the two, organic fertilizer can provide active carbon source for soil microorganisms, promote the metabolism and breeding of microorganisms. Biochar adsorbs and stores substances of different types and components in micropores, which provides nutrients for microbial communities, accelerates the accumulation of microbial necrosis, and enriches carbon in aggregates (Zhang et al., 2024). Due to the differences in physical, chemical and biological properties of different soil types, the experimental results in different regions are also significantly different, which leads to different responses of carbon components in soil aggregates to organic fertilizer and biochar.

The vegetable industry in southwest China is developed. The intensive cultivation and the cultivation mode of large water and large fertilizer, especially the excessive application of nitrogen fertilizer, accelerate the decomposition of soil organic carbon and lead to the deterioration of soil structure (Guo et al., 2020; Yang et al., 2022; Zhao et al., 2020;Zhu et al., 2021). In view of this, in order to solve the problem of soil organic carbon reduction in vegetable fields, this paper explored the relationship between the change of soil organic carbon components and aggregates and their stability, as well as the influence on carbon pool characteristics by applying organic fertilizer and biochar and their combined use. It is of great significance to optimize the management of vegetable soil carbon pool and the sustainable utilization of vegetable soil resources.

## 2 Materials and methods

### 2.1 Test site and materials

Planting types and planting and harvesting time of vegetables The positioning experiment began in 2019, and two crops of vegetables were planted alternately every year (Table 1).The experimental site is located in Chengjiang City, Yuxi City, Yunnan Province (N24 ° 39 ′ 6 ′′ E102 ° 54 ′ 48 ′′), Yunnan Agricultural University Agricultural Non-point Source Pollution Monitoring Station. The planting area belongs to the mid-subtropical plateau monsoon climate, with sufficient light, rain and heat in the same season, the annual average temperature is 11.9-17.5 °C, and the annual rainfall is 900-1200 mm. The typical vegetable field was selected, and the soil physical and chemical properties were as follows : pH 7.71, organic matter 19.88 g/kg, alkali-hydrolyzable nitrogen 79.03 mg/kg, available phosphorus 10.12 mg/kg, and available potassium 100.33 mg/kg. Biochar was prepared by pyrolysis of rice husk at 500 °C, N 0.35 %, P_2_O_5_ 0.72 %, K_2_O 2.43 %, pH 10.0, and the organic carbon content was 270.4 g/kg. Organic fertilizer is a commercial organic fertilizer (N 1.1 %, P_2_O_5_ 1.61 %, K_2_O 3.92 %) made from tobacco stem (leaf vein) as the main raw material. The pH is 7.6, and the organic carbon content is 218.5 g/kg, all of which are provided by Kunming Lishan Biotechnology Co., Ltd. Nutrient labeled amount of compound fertilizer was N-P_2_O_5_-K_2_O : 18-17-17.

| crops | species of vegetable | implantation time | harvest time |
| --- | --- | --- | --- |
| The first crop | lettuce | August 2019 | October 2019 |
| The second crop | lettuce | November 2019 | December 2019 |
| The third stubble | baby cabbage | May 2020 | July 2020 |
| The fourth crop | lettuce | September 2020 | October 2020 |
| The fifth crop | lettuce | May 2021 | June 2021 |
| The sixth crop | baby cabbage | August 2021 | October 2021 |
| The seventh crop | broc-coli. | April 2022 | July 2022 |
| The eighth crop | lettuce | August 2022 | September 2022 |
| The ninth crop | broc-coli. | March 2023 | June 2023 |
| The tenth crop | baby cabbage | September 2023 | November 2023 |

### 2.2 experimental design

The field randomized block design was adopted in the positioning test. Two crops of vegetables were treated during soil preparation each year :Single application of chemical fertilizer as control (CK) ; organic fertilizer completely replaces chemical fertilizer (OF, alternative application of total nutrients such as CK) ;Organic carbon such as biochar and organic fertilizer was applied, and nutrient deficiency was supplemented by chemical fertilizer (B) ; organic fertilizer and biochar 1/1, with OF, B treatment and other carbon inputs, insufficient nutrients supplemented by chemical fertilizer (BF).Each treatment set three replicates, a total of 12 experimental plots, each plot area of 27 m2, each plot interval of 20 cm. Biochar and organic fertilizer were fully mixed with 0-20cm soil (0.2m high, 2m wide) before each vegetable planting. 30 % of the compound fertilizer was mixed with the surface soil as the base fertilizer, and 70 % was used as the top dressing and watering. A total of 10 crops of vegetables were planted in 5 years, and local seedlings were purchased for transplanting. The fertilization schemes of different kinds of vegetables are shown in Table 2. Lettuce was applied twice after 10 days and 20 days of planting, 35 % each. The first topdressing of broccoli was 20 % at 20 days after transplanting, the second topdressing was 20 % at budding, and the third topdressing was 30 % at the stage of flower bulb expansion. The baby vegetables were applied four times at the seedling stage, rosette stage, early heading stage and late heading stage, and the fertilization method was watering. Pest prevention and weeding measures are carried out in accordance with local customs.

**Table 2.**
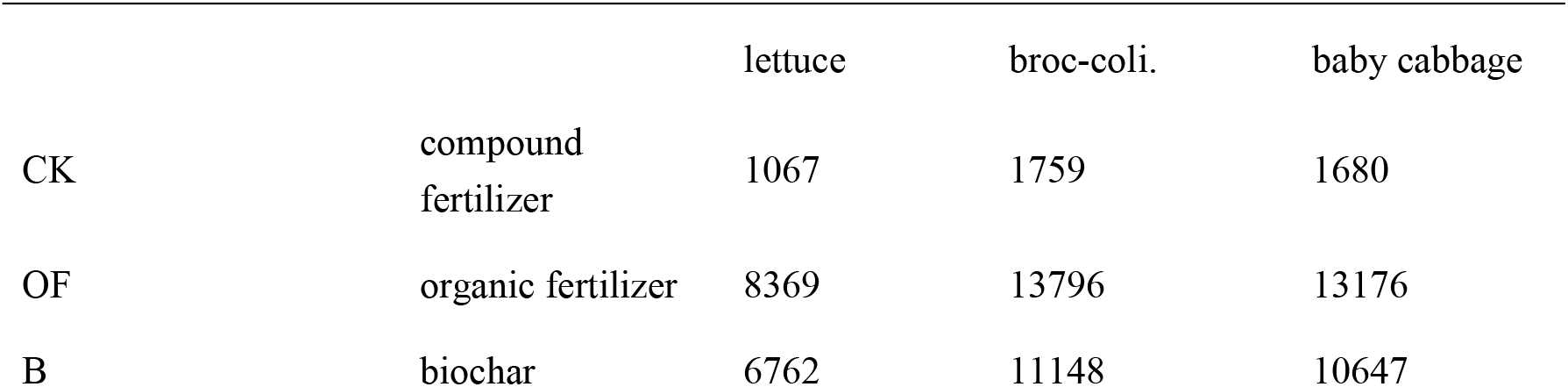
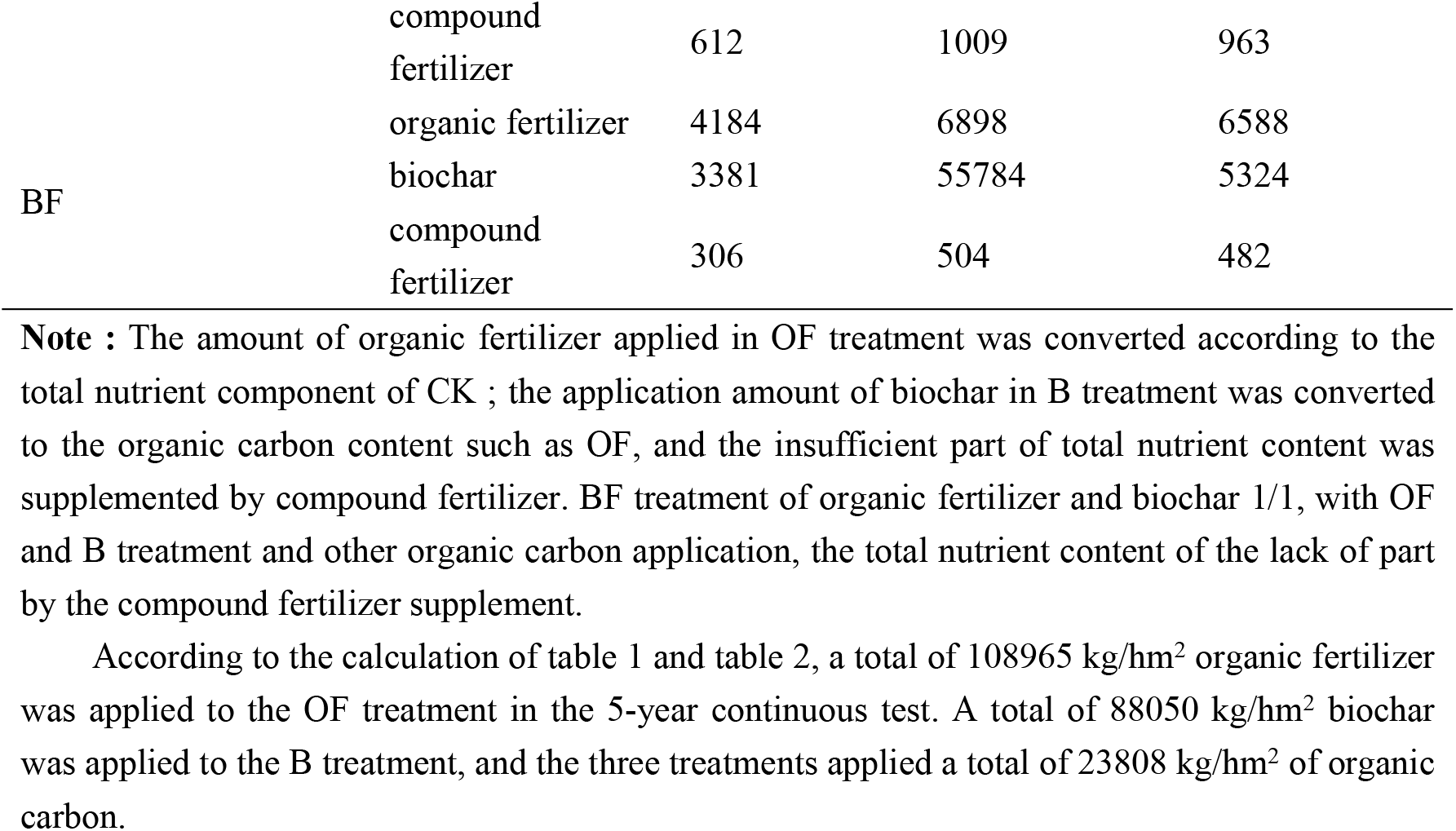
Fertilization amount of different vegetables per crop (kg/hm2)

According to the calculation of table 1 and table 2, a total of 108965 kg/hm2 organic fertilizer was applied to the OF treatment in the 5-year continuous test. A total of 88050 kg/hm2 biochar was applied to the B treatment, and the three treatments applied a total of 23808 kg/hm2 of organic carbon.

### 2.3 Sample collection and determination

From 2019 to 2023, in addition to measuring the yield of vegetables after each harvest, three mixed soil samples of 0-20 cm plough layer were collected. After air drying, the soil physical and chemical properties were determined by 20 mesh and 100 mesh sieves : alkali-hydrolyzable nitrogen, available phosphorus, available potassium, pH and other indicators. After the last harvest of vegetables in November 2023, five ring knife samples were taken from the plough layer of each treatment plot with stainless steel ring knife, and marked for the determination of soil bulk density. In each plot, the undisturbed soil samples of 0-20 cm soil layer were taken, and 5 points were fully mixed according to the ‘S ’ route. Three mixed samples were taken from each experimental plot. After removing gravel and plant residues, they were divided into 2 soil samples. One fresh sample was sealed and stored at 4 °C for screening of aggregates, and the particle size of aggregates was determined. One was used for the determination of soil organic carbon components. They are Soil Organic Carbon (SOC), Particulate Organic Carbon (POC), Readily Oxidizable Organic Carbon (ROC), Microbial Biomass Carbon (MBC), Dissolved Organic Carbon (DOC).One for the determination of soil organic carbon components, respectively, soil organic carbon (Soil Organic SOC with potassium dichromate external heating method; ROC with potassium permanganate oxidation method; MBC fumigation with chloroform; DOC was measured by TOC analyzer; POC was determined by sodium hexametaphosphate dispersion method. In addition, the calculation formula of soil inert organic carbon (IOC) is : IOC = SOC-ROC The calculation formula of the determination index is as follows :

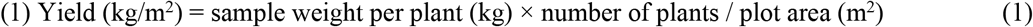

(2) Soil mechanical stability and water stability > 0.25mm aggregate content calculation formula:

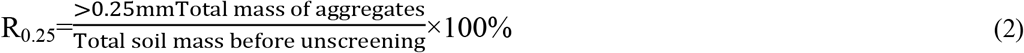

(3) The calculation formula of the destruction rate of soil aggregate structure :

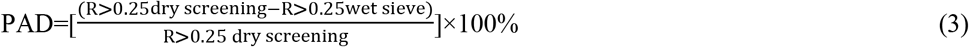

(4) The calculation formula of average mass diameter and geometric mean diameter of aggregates :

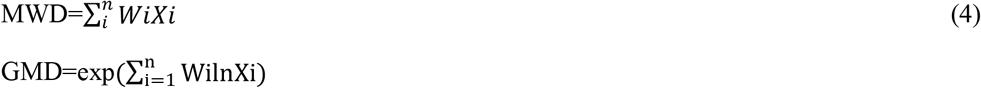

In the formula, n, Xi and Wi are the number of particle size groups, the average diameter of the particle size aggregates (mm), and the percentage of the mass of the particle size aggregates to the soil mass (%).

(5) Fractal dimension D :

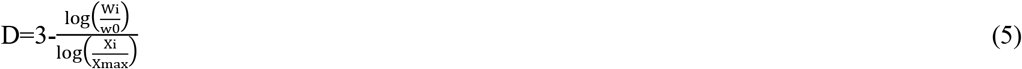

Wi, W0, Xi and X max are the accumulated mass of soil particles with diameter less than 0.05 mm, g ; the sum of all grain-size soil mass, g ; the average particle size of aggregates in the range of i particle size, mm ; the average diameter of the largest grain size of soil particles, mm.

(6) Calculation of soil organic carbon storage:

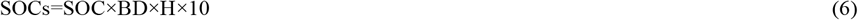

In the formula : SOCs, SOC, BD, H, respectively, organic carbon storage (t/hm2), soil organic carbon content (g/kg), bulk density (g/cm3), soil thickness (0.2m).

(7) Soil carbon pool management index:

Carbon pool index (CPI) = sample total carbon A content (g/kg) / original soil total carbon content (g/kg) ; carbon pool activity (A) = active carbon content (g/kg) / inactive carbon content (g/kg) ; carbon pool activity index (AI) = sample carbon pool activity / original soil carbon pool activity ; carbon pool management index (CPMI) = carbon pool index × carbon pool activity index × 100 = CPI × AI × 100

### 2.4 Data analysis

Microsoft Excel 2021 was used to collate and summarize the original data, and SPSS.26 was used to analyze the variance of the data. The least significant difference method (Duncan) was used to test the significance of the difference (P < 0.05). Origin 2022 mapping and Canoco 5.0 were used for redundancy analysis.

## 3 Results and analysis

### 3.1 Changes of soil organic carbon fractions

As shown in fig 1, compared with the control (CK), the contents of organic carbon (SOC), particulate organic carbon (POC), readily oxidizable organic carbon (ROC), soluble organic carbon (DOC), microbial biomass carbon (MBC) and inert organic carbon (IOC) in biochar (B), organic fertilizer (OF) and their combined application (BF) were significantly increased. Among them, SOC, POC, IOC, B treatment was better than OF treatment, and showed significant differences, while ROC, DOC and MBC, OF treatment was better than B treatment, and the difference was significant.

**Fig. 1.**
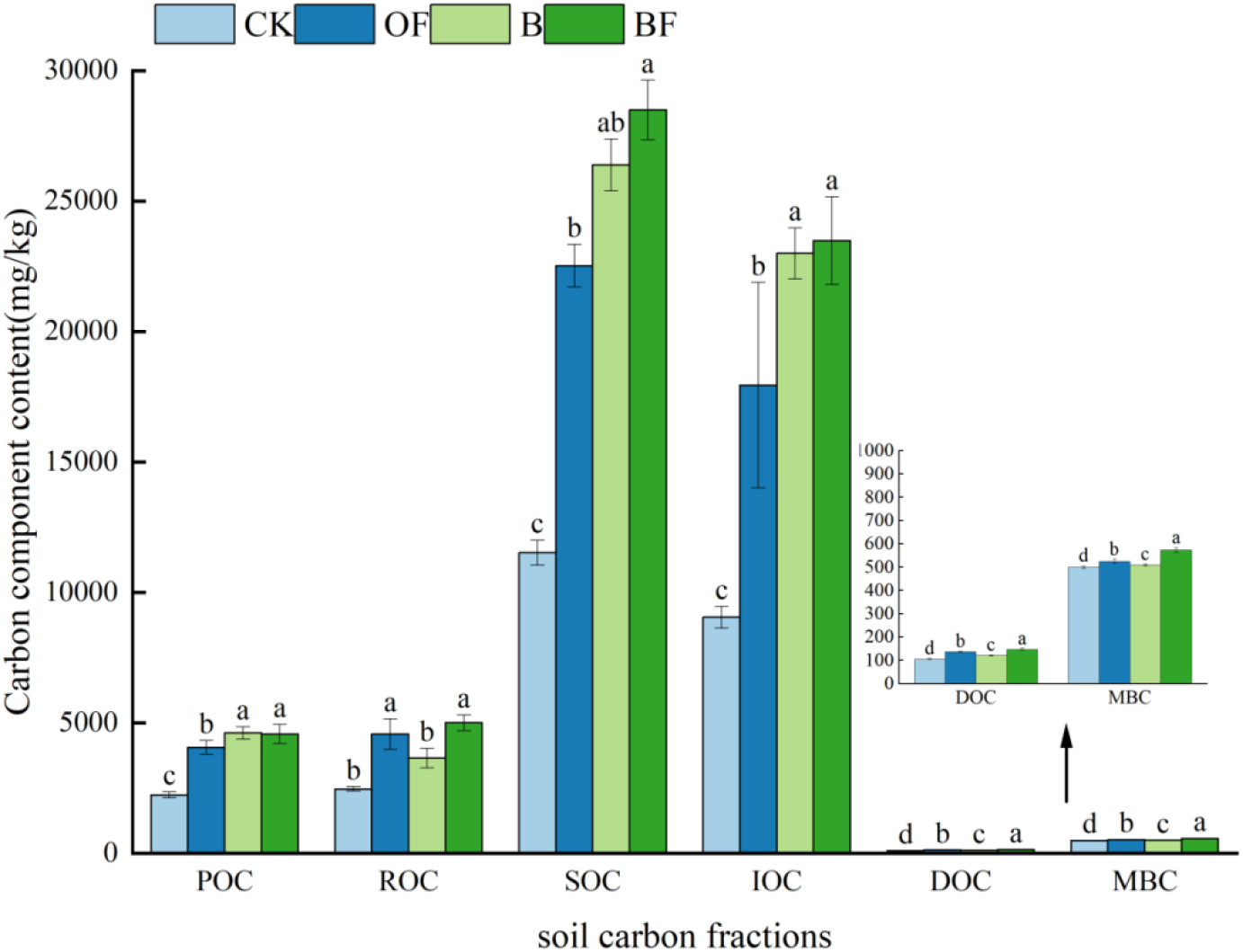
Organic carbon fractions in soil under different fertilization regimes.

### 3.2 Effects of different carbon sources on the distribution and stability parameters of soil aggregates

Fig.2 is the distribution of soil aggregates under different fertilization treatments. Compared with CK, the soil water-stable aggregates with particle size < 0.25 mm in OF, B and BF treatments were significantly reduced by 10%, 3% and 10%, respectively. In Table 3, compared with CK, R0.25 was significantly increased under OF and BF treatments, which increased by 10.51% and 10.7 %, respectively, and the effect under BF treatment was the best. Compared with CK, the average mass diameter (MWD) of aggregates under OF and BF treatments was significantly increased by 17.0% and 15.0%, respectively. The geometric mean diameter (GMD) increased significantly under OF, B and BF treatments, increasing by 11.0%, 3.0% and 10.0%, respectively. The data showed that the application of organic fertilizer (OF) had the best effect on increasing the MWD and GMD of soil aggregates. This is also true in reducing the soil aggregate structure destruction rate (PAD%) and fractal dimension (D).

**Table 3.** Effects of different fertilization treatments on soil aggregate stability indexes.

| treatment | water stability |  |  |  |  |
| --- | --- | --- | --- | --- | --- |
|  | MWD | GMD | R0.25% | PAD% | D |
| CK | 0.53±0.09b | 0.27±0.01c | 39.72±3.7b | 57±2.1a | 2.25±0.22a |
| OF | 0.70±0.01a | 0.38±0.002a | 50.23±0.6a | 47.66±0.34b | 1.77±0.05b |
| B | 0.55±0.01b | 0.30±0.002a | 43.29±0.1b | 54.22±0.36ab | 2.22±0.02a |
| BF | 0.68±0.02a | 0.37±0.02a | 50.42±1.9a | 48.04±2.35b | 1.94±0.14b |

**Fig. 2.**
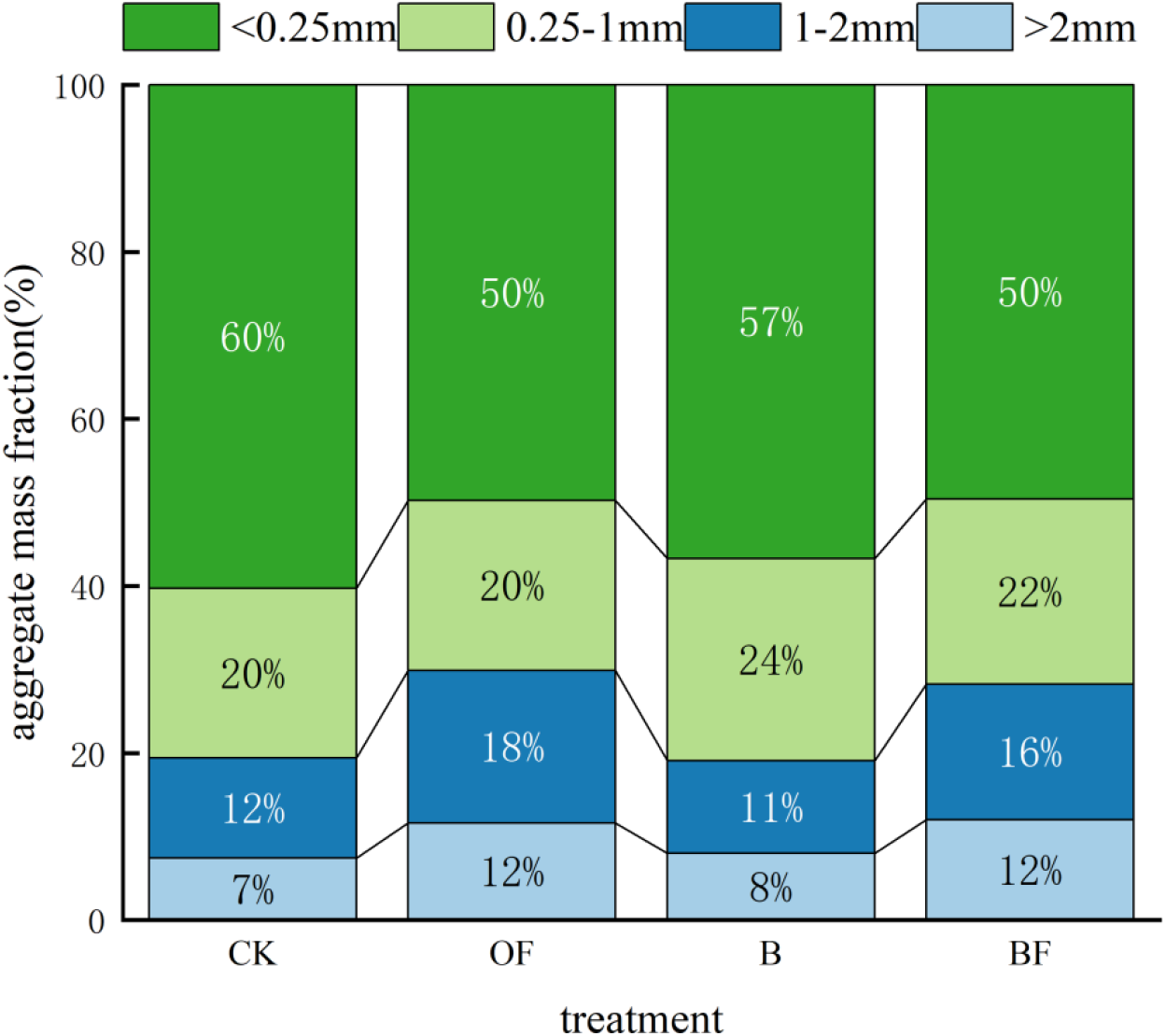
Stable soil water aggregates under different fertilization treatments.

### 3.3 Effects of different carbon sources on the distribution ratio of each carbon component in different size aggregates

After separating the aggregates of each particle size, the distribution of carbon components in the aggregates of each particle size was analyzed (Fig.3).The distribution ratio of R0.25 in different carbon fractions was increased under OF, B and BF treatments. The proportion of R0.25 in aggregates under the four treatments in DOC was BF > CK > OF > B ; The distribution ratio of R0.25 in MBC was BF = B = OF = CK ; the distribution ratio of R0.25 in POC was B > CK > OF > BF ; the distribution ratio of R0.25 in EOC was CK = OF > BF > B.

**Fig. 3.**
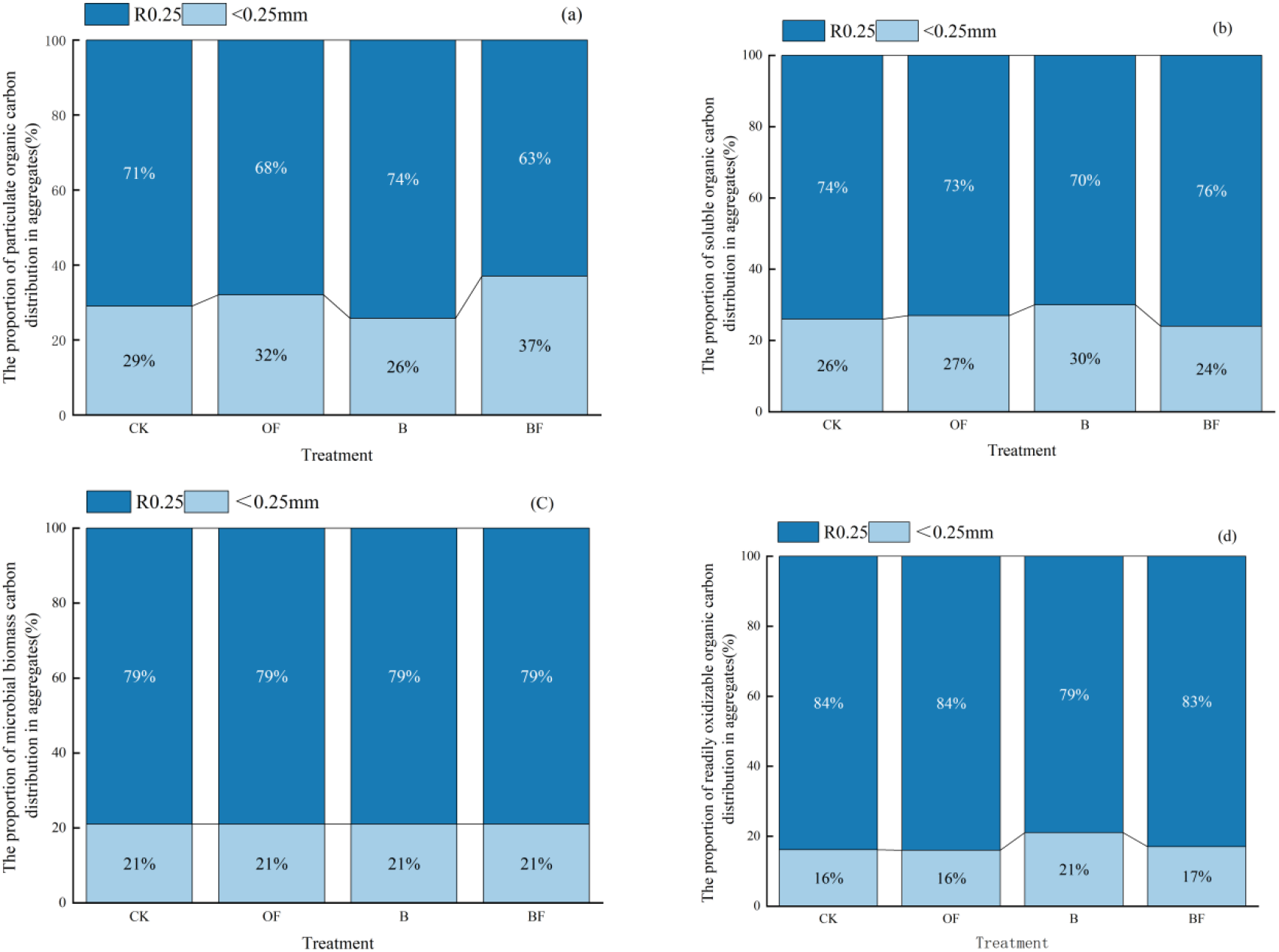
(a) The proportion of particulate organic carbon distribution in aggregates (%); (b) The proportion of soluble organic carbon distribution in aggregates; (c) The proportion of microbial biomass carbon distribution in aggregates; (d) The proportion of readily oxidizable organic carbon distribution in aggregates

### 3.4 Correlation analysis between soil organic carbon fractions and aggregates and their stability

The correlation analysis between soil aggregates and their stability and organic carbon components is shown in Fig.4.In Fig.4a, the content of soil aggregates with particle size > 0.25mm was positively correlated with SOC, POC, ROC, IOC, MBC and DOC, among which the soil aggregates with particle size > 2mm were significantly positively correlated with MBC and DOC. Among soil aggregates and aggregate stability, MWD, GMD and R0.25 were significantly positively correlated with soil aggregates with particle size > 1mm, and significantly negatively correlated with soil aggregates with particle size > 0.25mm.There was a significant negative correlation between D and soil aggregates with particle size > 1mm, and a significant positive correlation with soil aggregates with particle size > 0.25mm.Between soil carbon fractions and aggregate stability, each carbon fraction was negatively correlated with PAD and D, and positively correlated with MWD, GMD and R0.25.In Fig.4b, the content of soil aggregates with particle size > 2mm was significantly positively correlated with MWD, GMD and R0.25, and significantly negatively correlated with PAD and D ; the content of 1-2mm soil aggregates was significantly positively correlated with MWD, GMD and R0.25, and significantly negatively correlated with PAD and D ; the content of soil aggregates < 0.25 mm was significantly negatively correlated with MWD, GMD and R0.25, and significantly positively correlated with PAD and D.

**Fig. 4.**
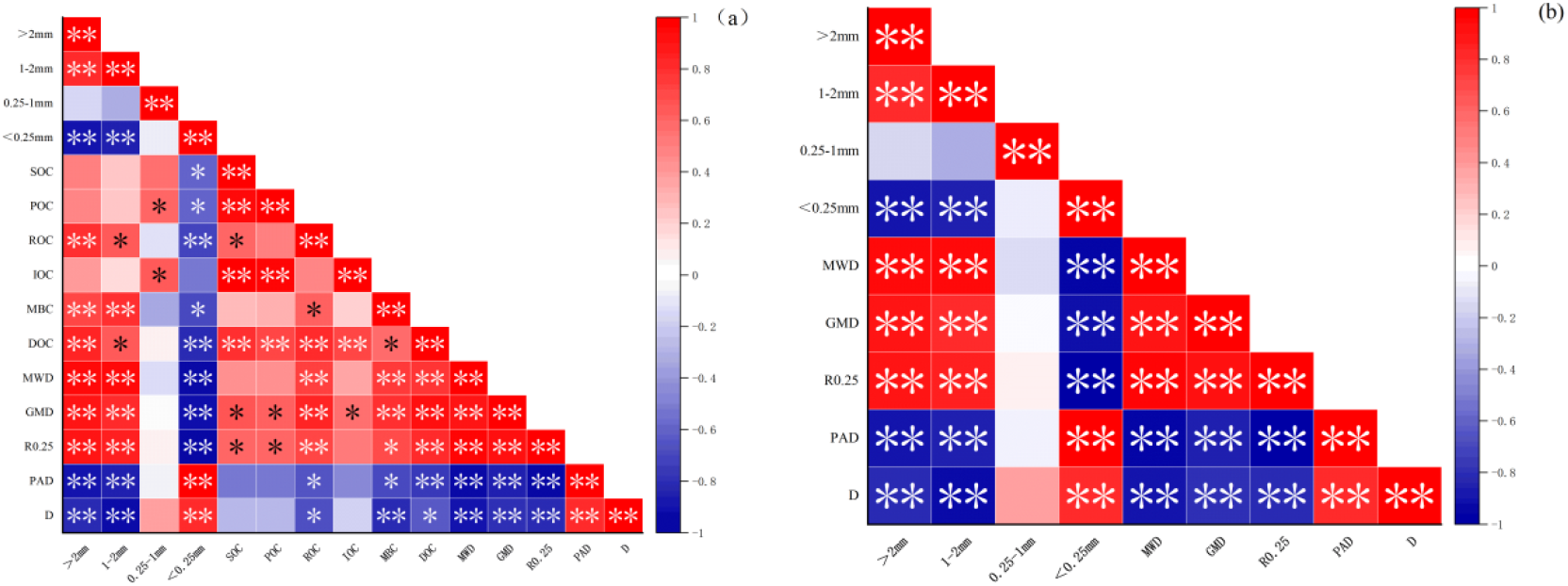
(a) correlation analysis of carbon composition and stability of aggregates; (b) correlation analysis of aggregate mass fraction and its stability

### 3.5 Effects of continuous application of organic fertilizer and biochar on vegetable yield

Fig.5 shows the vegetable yield of continuous application of biochar and organic fertilizer for 5 years. The organic substitution (OF) significantly reduced the vegetable yield compared with the control, and the application of biochar (B) and the combined application (BF) increased the vegetable yield. In general, the order was BF treatment > B treatment > CK > OF treatment.

**Fig. 5.**
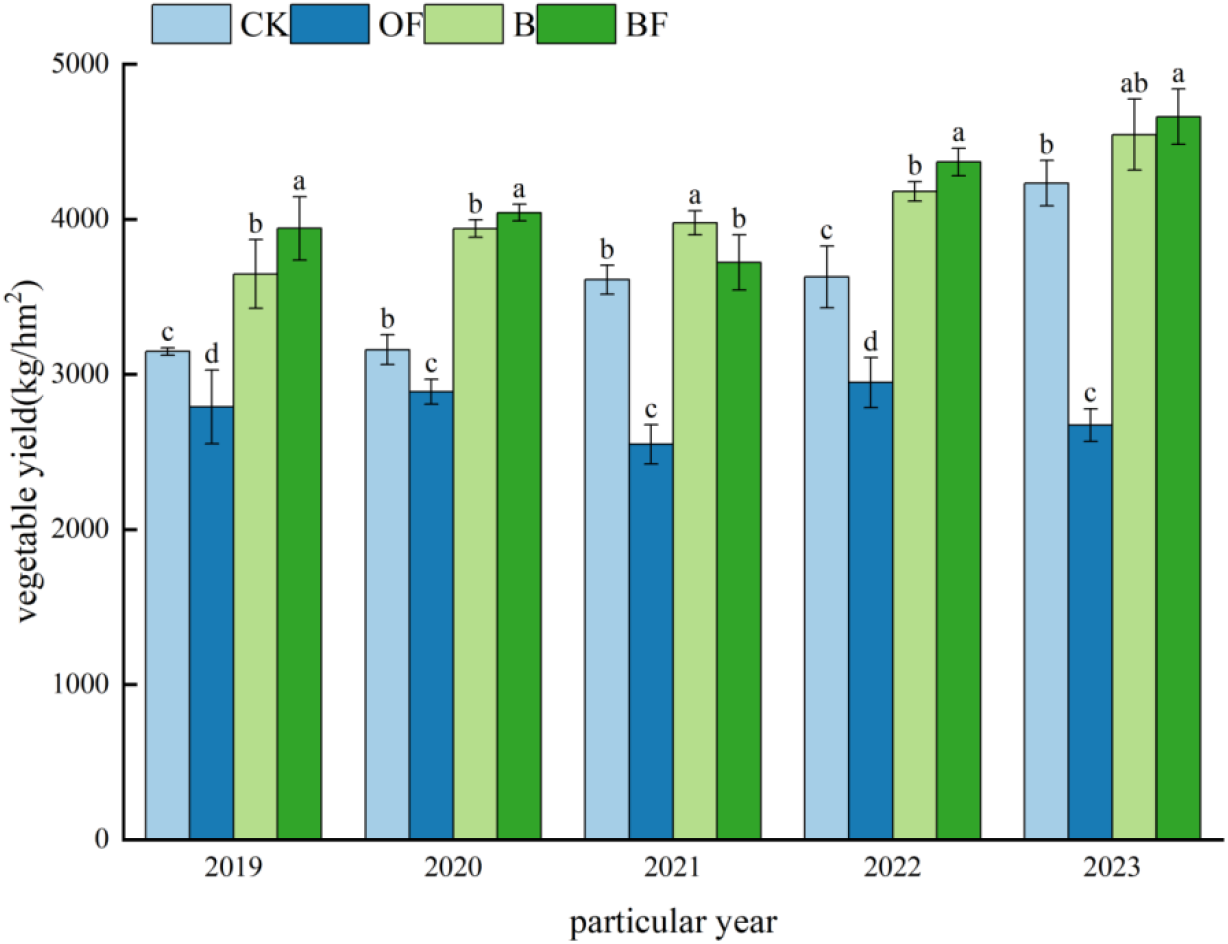
Vegetable yield under different treatments

### 7.6 Effects of continuous application of organic fertilizer and biochar on the dynamic changes of soil conventional five items

As shown in a b c of Fig.6, the contents of alkali-hydrolyzable nitrogen, available phosphorus and available potassium in soil decreased first and then increased. By 2023, the contents of alkali-hydrolyzable nitrogen, available phosphorus, available potassium and orga nic matter in BF treatment were better than those in B and OF treatment, while the increase of available phosphorus content in OF treatment was better than that in B treatment, and the contents of alkali-hydrolyzable nitrogen, available potassium and organic matter in B treatment were better than those in OF treatment. Soil pH, compared with CK treatment, the other three treatments significantly increased pH. From 2021, the pH under OF, B, and BF treatments gradually increased. By 2023, each treatment showed B > BF > OF > CK.

**Fig. 6.**
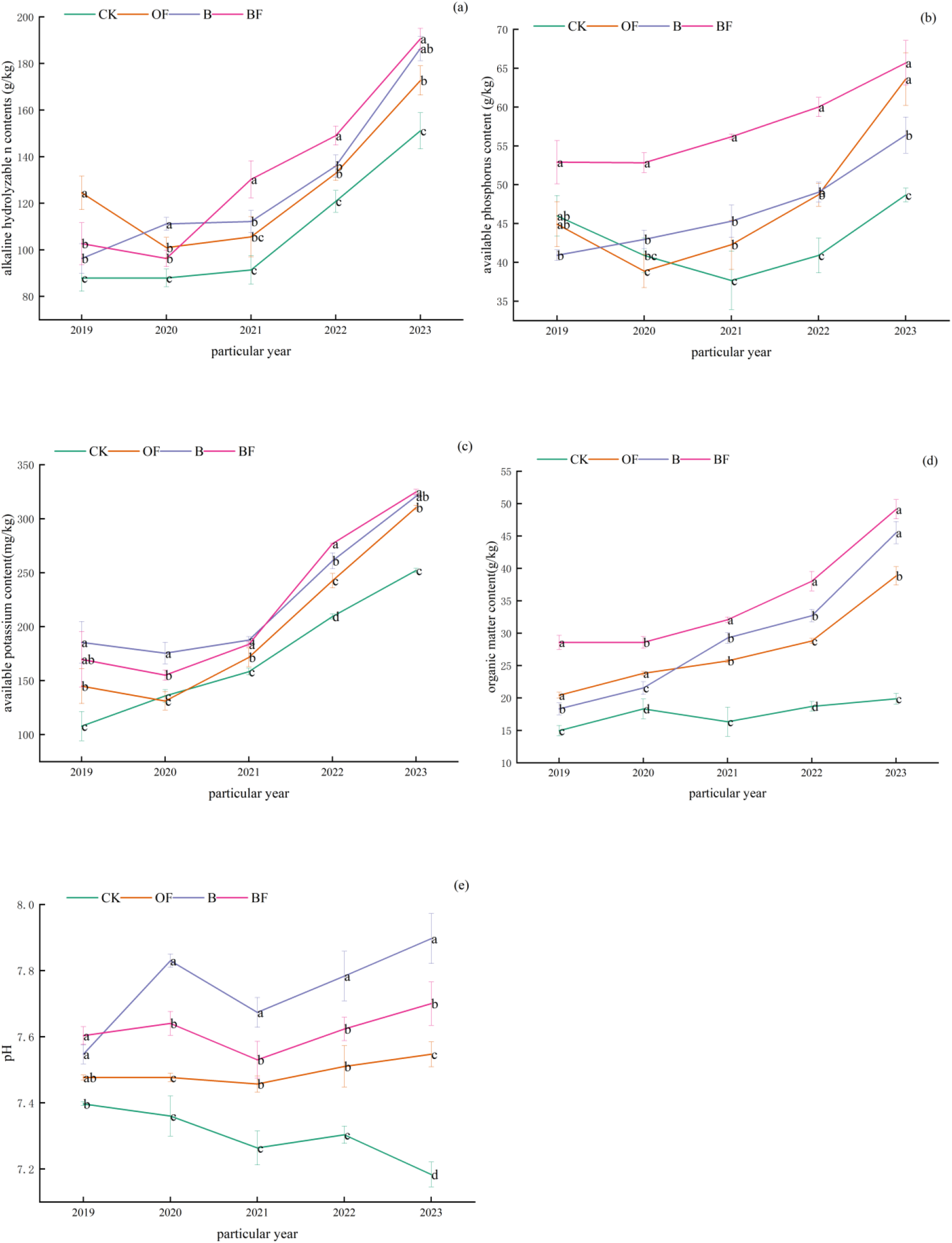
Dynamic changes of soil available nutrients (a) alkaline hydrolysable n contents(g/ kg);(b)available phosphorus content (g/kg); (c) available potassium content (mg/kg); (d) org anic matter content (g/kg); (e) soil pH.

### 3.7 Correlation between soil aggregates, organic carbon fractions and physical and chemical properties

As shown in figure 7, the moisture content was significantly positively correlated with AK and AP, and negatively correlated with PAD and D ; AP was significantly positively correlated with R0.25 and significantly negatively correlated with D ; AK was positively correlated with R0.25 and negatively correlated with PAD. Soil moisture content was positively correlated with TOC, POC, DOC, ROC and IOC. pH, AN and AP were significantly positively correlated with POC and ROC, and significantly positively correlated with MBC and DOC. AK was significantly positively correlated with POC, ROC, IOC and DOC.

**Fig. 7.**
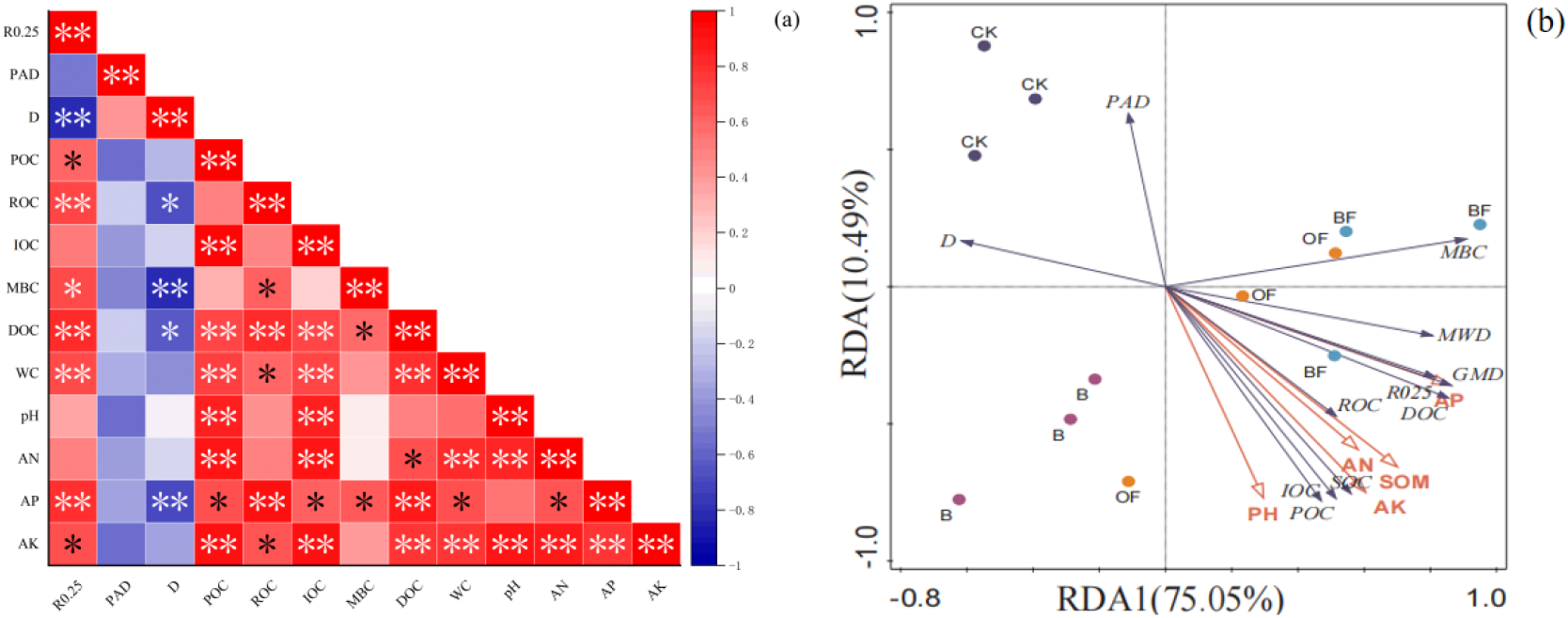
(a) Correlation between aggregate stability and soil physical and chemical properties (b) Redundancy analysis of aggregate stability and soil physical and chemical properties

Correlation analysis showed that there was a strong correlation between soil physical and chemical properties and soil aggregates and organic carbon components. Therefore, redundancy analysis was carried out with soil physical and chemical properties as environmental variables and other indicators as response variables. The results could explain 85.54%. It can be seen from the figure that pH, AN, SOM, AK and AP were positively correlated with ROC, POC, DOC, MBC, IOC, TOC, R0.25, MWD and GMD, indicating that the increase of available nutrient content in soil contributed to the increase of soil organic carbon components. The pH, AN, SOM, AK and AP were negatively correlated with PAD and D, and AP and SOM were the main factors affecting the stability and carbon components of aggregates.

### 2.8 Effects of different fertilization treatments on the stability of soil organic carbon pool

Table 4 shows the stability of soil organic carbon pool under different fertilization treatments. Compared with CK treatment, soil organic carbon storage increased significantly under OF, B and BF treatments, which increased by 29.73, 36.44 and 39.521000 kg/hm2, respectively. B increased by 6.711000 kg/hm2 compared with OF treatment, and the effect of combined application of the two was the best. BF increased by 9.791000 kg/hm2 compared with OF treatment. The variation range of carbon pool activity index AI was 1.13 ∼ 1.56. Compared with CK treatment, AI was significantly increased under BF treatment. The carbon pool index ranged from 1.26 to 1.96. Compared with CK treatment, the CPI under OF, B and BF treatments was significantly increased by 39.00%, 68.00% and 70.00%, respectively. It can be seen that when the CPI is increased, the effect of organic fertilizer and biochar is the best. The carbon pool management index ranged from 141.55 to 303.10. Compared with CK treatment, CPMI under OF, B and BF treatments was significantly increased by 140.49%, 66.29% and 161.55%, respectively. Among them, CPMI under B treatment was significantly lower than that under OF treatment, which was lower than that under OF treatment by 74.20%.

**Table 4.** Effects of different fertilization treatments on soil organic carbon pool characteristics.

| Treatment | total soil organic carbon<br>(1000kg/hm<sup>2</sup>) | AI | CPI | CPMI |
| --- | --- | --- | --- | --- |
| CK | 32.89±1.37c | 1.13±0.11b | 1.26±0.11c | 141.55±2.02d |
| OF | 62.62±2.84b | 1.40±0.07ab | 1.65±0.08b | 282.04±2.16b |
| B | 69.33±2.60ab | 1.20±0.18b | 1.94±0.11a | 207.84±4.90c |
| BF | 72.41±2.74a | 1.56±0.23a | 1.96±0.13a | 303.10±8.32a |

### 3.9 Comprehensive evaluation of different fertilization treatments on carbon pool characteristics and soil fertility in vegetable field

Principal component analysis was carried out on soil organic carbon components, soil aggregates, organic carbon storage, carbon pool activity index, carbon pool index, carbon pool management index and soil routine five items, and the principal components were extracted according to the standard of eigenvalue greater than 1. The results showed that there were two principal components affecting the stability of soil carbon pool, as shown in Table 6. The cumulative variance contribution rate was 90.24%, which had good representativeness and could be used to evaluate the effects of different fertilization treatments on the stability of soil carbon pool.

**Table 5.** Coefficient and contribution rate of each index.

| index | Principal component 1 | Principal component 2 |
| --- | --- | --- |
| POC | 0.89 | 0.39 |
| DOC | 0.93 | -0.18 |
| MBC | 0.58 | -0.63 |
| ROC | 0.81 | -0.36 |
| IOC | 0.86 | 0.48 |
| R<sub>0.25</sub> | 0.83 | -0.41 |
| <0.25mm | -0.83 | 0.41 |
| total soil organic carbon | 0.93 | 0.34 |
| AI | 0.59 | -0.77 |
| CPI | 0.78 | 0.56 |
| CPMI | 0.94 | -0.29 |
| pH | 0.75 | 0.55 |
| AN | 0.83 | 0.45 |
| AP | 0.92 | -0.28 |
| AK | 0.95 | 0.26 |
| SOM | 0.97 | -0.05 |
| eigenvalue | 11.41 | 3.03 |
| variance contribution rate% | 47.53 | 42.71 |
| Cumulative contribution rate% | 47.53 | 90.24 |

**Table 6.**
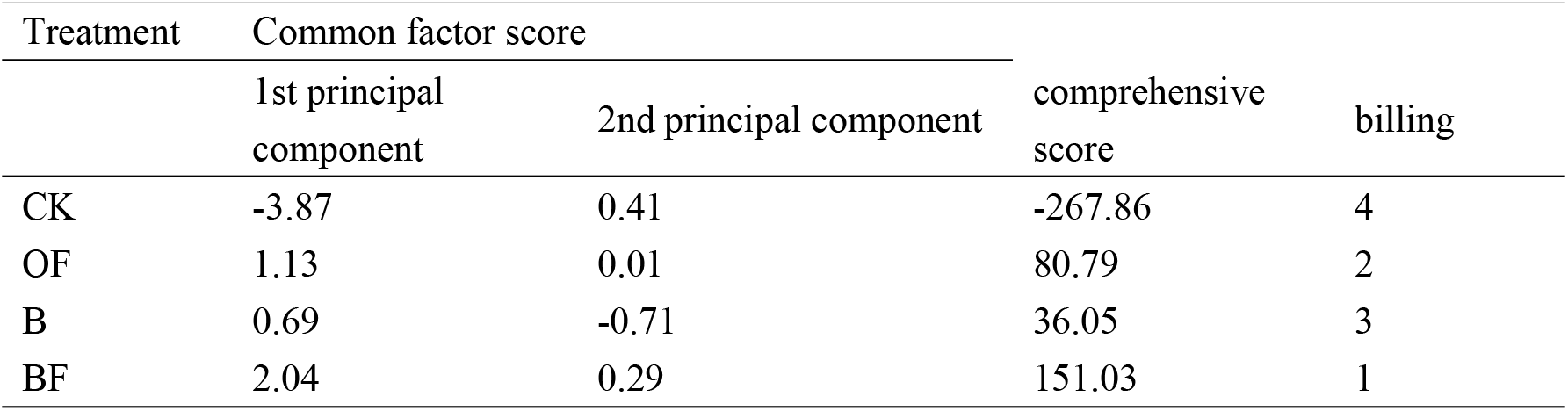
Common factor scores and comprehensive scores of different fertilization methods.

The first two principal components are selected, and the obtained feature vector is multiplied by the standardized data to obtain the principal component expression. According to the proportion of the eigenvalues corresponding to the principal component to the sum of the total eigenvalues of the extracted principal components, the principal component synthesis model is obtained. According to the principal component synthesis model, the comprehensive principal component value is calculated and sorted according to the comprehensive principal component value. As shown in Table 7, the comprehensive scores of each treatment were ranked as BF treatment > OF treatment > B treatment > CK treatment, indicating that the combined application of organic fertilizer and biochar had the highest comprehensive evaluation in improving the stability of soil organic carbon pool and soil fertility.

## 4 Discussion

### 4.1 Effects of organic manure and biochar on soil carbon fractions, soil aggregates and stability

The proportion of water-stable soil aggregates, mean weight diameter, geometric mean diameter, aggregate destruction rate and fractal dimension of R0.25 are all important indicators for evaluating the degree of soil agglomeration (Tian et al., 2020).This study found that organic substitution and application of biochar can significantly increase the content, geometric mean diameter and mean weight diameter of soil macroaggregates, and reduce aggregate destruction rate and fractal dimension (Yang et al., 2023).The possible reason is that organic substitution accelerates the process of soil humification and promotes the cementation of aggregates. Biochar promotes the formation of aggregates by increasing organic matter, and because of the huge specific surface area and developed pore structure of biochar, it provides a rich living space for microorganisms and accelerates the accumulation of microbial necrosis to form stable aggregates. (Peng et al., 2015) Among them, the effect of biochar is better than that of organic fertilizer, which may be due to the low microbial activity of aggregates mediated by biochar, which is not easy to be decomposed, while organic fertilizer is mainly formed by cementation. Aggregates have strong microbial activity. Therefore, its ability to improve the stability of aggregates is lower than that of biochar (P. Xu et al., 2024).Although both organic fertilizer and biochar can promote the formation of aggregates, the combination of biochar and organic fertilizer can best increase the content of soil macroaggregates in this study. The possible reason is that the two coordinate with each other to change the microbial community structure, accelerate the humification process, promote microbial metabolism and clay cementation of the bacteria themselves, and increase the content of macroaggregates and the stability of soil structure, which is consistent with the research conclusions of Song et al (Zhang et al., 2022).Correlation analysis showed that the particle size of aggregates was a significant factor affecting the stability of aggregates. The larger the aggregates, the stronger the stability. However, some studies have found that the higher the organic carbon content, the stronger the stability of aggregates.

### 4.2 Effects of organic fertilizer and biochar on vegetable yield and soil conventional five items

Vegetable yield is closely related to its genetic characteristics, soil type, nutrient status and climatic conditions. The supply of soil nutrients is one of the most important factors affecting yield (Chandra & Bhattacharya, 2019).Organic fertilizer instead of chemical fertilizer treatment and single application of biochar and organic fertilizer combined with biochar treatment can significantly increase vegetable yield (Li et al., 2023).This study found that the yield of vegetables was significantly reduced under the treatment of organic fertilizer completely replacing chemical fertilizer, while the yield was significantly increased under the treatment of single biochar and organic fertilizer combined with biochar. The slow release of organic fertilizer nutrients and the lack of coordination with the nutrients required for vegetable growth are the main reasons for the reduction of vegetable yield. The application of biochar increased vegetable yield, mainly due to the adsorption of nutrients by biochar, which increased the supply of nutrients required by plants (Sun et al., 2022).The reason why the combined application of organic fertilizer and biochar can increase the yield of vegetables is that the interaction between biochar and organic fertilizer changes the soil physical and chemical conditions and microbial flora, thus improving the nutrient supply capacity (Velli et al., 2021).There are also some reports that organic cultivation can reduce crop yield, depending on the fertilizer requirement characteristics of different crops and the length of the growth period(Feng et al., 2023).Organic fertilizer completely replaces chemical fertilizer, that is, organic cultivation conditions can ensure crop yield. Under the condition of this experiment, the yield of organic fertilizer completely replacing chemical fertilizer was lower than that of the control chemical fertilizer treatment in 10 stubble experiments. The possible reason was that the growth period of leafy vegetables was shorter, and the organic fertilizer did not decompose and release enough nutrients during this period. In the organic substitution experiment of tomato and cucumber, our research group found that the biomass decreased in the early stage of growth, but it surpassed the single chemical fertilizer treatment in the later stage of yield formation. Therefore, the complete substitution of organic fertilizer for chemical fertilizer on crops with short growth period may reduce crop yield.

Soil available nutrients are important indicators to measure soil fertility. Organic fertilizer content, alkali-hydrolyzable nitrogen content, available phosphorus content, available potassium content and pH reflect soil nutrient status to a certain extent (Zhan et al., 2017).In this study, each treatment significantly increased the content of soil available nutrients, and changed significantly with time, showing BF treatment > B treatment > F treatment > CK. This is because with the input of organic carbon, the function of soil microorganisms was changed and the fixation of nutrients by microorganisms was accelerated (Chaopricha & Marín-Spiotta, 2014).In addition, since the input of organic carbon also accelerates the formation of soil aggregates, different nutrients are adsorbed inside the aggregates and further form large aggregates through cementation and other effects. However, different carbon sources have different effects on the formation of aggregates [35](Li et al., 2024), which may lead to different distribution of carbon sources and nutrients inside the aggregates, thereby affecting soil microecology. In this study, in general, the effect of biochar is better than that of organic fertilizer. This may be because biochar is a carbon-containing material formed by thermal cracking, which has a huge specific surface area and cation exchange capacity, and has an adsorption effect on nutrients.

### 4.3 Effects of organic fertilizer and biochar on carbon fractions in soil aggregates

DOC and EOC are important indicators of soil quality, and the increase in POC helps to improve soil quality(Wan et al., 2024). In this study, the application of organic fertilizer and biochar changed the content of carbon components in aggregates with different particle sizes. Among them, BF significantly increased the content of POC and ROC in macroaggregates, while organic substitution and biochar application decreased the content of DOC in macroaggregates, which was basically consistent with the research conclusions of (Wang et al., 2024). The reason is that the input of exogenous carbon source will increase the excitation effect of soil and accelerate the mineralization and decomposition of soil organic carbon. However, different carbon sources have different abilities to promote mineralization and decomposition. Stable carbon can promote the decomposition of organic carbon. In this study, the interaction between organic fertilizer and biochar improves soil function and enhances carbon sequestration. Organic substitution accelerates the consumption of organic carbon due to the high activity of microorganisms and reduces the sequestration of organic carbon. The application of biochar accelerates the decomposition of soil organic carbon and changes the content of POC and DOC to a certain extent (Liu et al2023). found that the application of organic fertilizer and biochar could significantly increase the content of easily oxidizable organic carbon, while Chen et al. ’ s research results showed that the application of organic fertilizer and biochar could reduce the content of easily oxidizable organic carbon. The contradictory conclusion may be due to the different cultivated crops, soil types and climatic conditions. In this study, the application of organic fertilizer and biochar reduced the content of easily oxidized organic carbon, because the application of organic fertilizer and biochar stimulated the activity of microorganisms, accelerated the reproduction of microorganisms and the consumption of easily oxidized organic carbon. The correlation analysis showed that the particle size of aggregates would affect the soil carbon components, which was similar to the conclusion of (He et al2024). In summary, aggregates promoted the sequestration of organic carbon, but due to different carbon source inputs, the internal carbon components of different aggregates were also significantly different.

### 4.4 Effects of organic fertilizer and biochar on carbon storage and carbon pool characteristics

The application of organic fertilizer and biochar can change carbon components and increase soil carbon sequestration. In this study, the application of organic fertilizer and biochar significantly increased soil organic carbon content and increased carbon storage. The effect of biochar application was better than that of organic fertilizer, which was consistent with the research results of (Zhang et al2024). Exogenous carbon input is a measure to increase soil organic carbon, and the application of organic fertilizer and biochar can increase carbon storage. However, organic fertilizer is mainly rich in active organic carbon microorganisms, while biochar is mainly rich in stable organic carbon, which is not easy to be utilized by microorganisms. Therefore, the effect of biochar on improving organic carbon storage is better than that of organic fertilizer. Soil carbon pool management index and carbon pool activity index can sensitively reflect the dynamic changes of soil carbon(Liu et al., 2024) .Previous studies have shown that carbon pool management index and activity index are closely related to tillage measures and fertilization methods(Wang et al., 2020) .In this study, compared with single application of chemical fertilizer, the application of organic fertilizer and biochar increased the carbon pool management index and activity index, and the effect of organic fertilizer on carbon pool management index and activity index was better than that of biochar. This may be because organic fertilizer is mainly rich in active organic carbon and has high microbial utilization, while the combination of organic fertilizer and biochar has the best effect on improving carbon pool management index and activity index. This is because the application of organic fertilizer and biochar changed the soil carbon composition and increased the content of soil total organic carbon and active organic carbon(Huanhuan et al., 2022), while the carbon pool management index was closely related to soil total organic carbon and easily oxidized organic carbon, and increased with the increase of soil organic carbon (P. Xu et al., 2024).In addition, the interaction between biochar and organic fertilizer improved the soil microenvironment and increased microbial activity, so the carbon pool management index and activity index were significantly improved. Biochar is superior to organic fertilizer in improving carbon pool index, which is mainly related to the properties of biochar and organic fertilizer. Biochar contains many aromatic functional groups and is not easy to decompose, while organic fertilizer is mainly rich in active organic carbon such as glucose, and microbial utilization is large. The degree of mineralization and decomposition of organic fertilizer is much higher than that of biochar. Therefore, the ability of biochar to improve carbon pool index is significantly better than that of organic fertilizer, but the combined application of biochar and organic fertilizer has the best effect. This is mainly because the combined application of biochar and organic fertilizer significantly changes microbial activity, increases microbial necrosis, agglomeration and cementation, and promotes the increase of soil carbon storage by increasing aggregates(P. D. Xu et al., 2024). Therefore, we can see. Because biochar and organic fertilizer provide different carbon sources, soil carbon pool stability and carbon pool activity have different responses to different carbon sources. Active organic carbon enhances carbon pool activity, while inert organic carbon enhances carbon pool index, and the combination of the two has the best effect on increasing carbon storage. From the perspective of conducive to the conservation and fertilization of cultivated land, the combined application of biochar and organic fertilizer has the best effect on improving cultivated land fertility and soil carbon pool (Zhang et al., 2024). Principal component analysis was used to comprehensively evaluate each index, and the effects of different fertilization treatments on carbon pool characteristics and soil fertility were comprehensively analyzed. The results showed that compared with conventional fertilization, the application of biochar and organic fertilizer could improve soil carbon pool characteristics and soil fertility. The effect of organic fertilizer was better than that of biochar, and the combined application of the two had the best effect, which was consistent with the results of previous studies(Li et al., 2024).

## 5 Conclusion

The application of biochar and organic fertilizer reorganized soil organic carbon from different components. Organic fertilizer improved the characteristics of carbon pool through the formation and development of large aggregates mediated by DOC and MBC, enhanced the aggregation ability of organic carbon components in aggregates, and promoted soil carbon sequestration in vegetable fields. Biochar itself is stable carbon and directly constitutes carbon sequestration. The combination of organic fertilizer and biochar has the best effect on soil fertilization and carbon sequestration.

## author contribution

Yuan Yang Peng : Write-original draft, visualization, validation, methodology, survey, formal analysis, data collation, conceptualization.

Jiang Ling Huang : Write-review and edit, supervise, methodology, data collation, conceptualization.

Yan Qin Peng: Write-review and edit, supervise, methodology, data collation, conceptualization. Yin Peng: Write-review and edit, supervise, methodology, data collation, conceptualization.

Shi Xiong Li: Write-review and edit, supervise, methodology, data collation, conceptualization.

Xi La Tu DaBu: Review and editing, supervision, project management, investigation, formal analysis, conceptualization.

## Declarations

### Open Access

This article is licensed under a Creative Commons Attribution 4.0 International License, which permits use, sharing, adaptation, distribution and reproduction in any medium or format, as long as you give appropriate credit to the original author(s) and the source, provide a link to the Creative Commons licence, and indicate if changes were made. The images or other third party material in this article are included in the article’s Creative Commons licence, unless indicated otherwise in a credit line to the material. If material is not included in the article’s Creative Commons licence and your intended use is not permitted by statutory regulation or exceeds the permitted use, you will need to obtain permission directly from the copyright holder.

The authors declare that they have no known competing financial interests or personal relationships that could have appeared to influence the work reported in this paper.

## Acknowledgements

We are grateful to the National Natural Science Foundation of China (41967015) and Fu xian Lake Ecological Environment Protection Project (ZC530400201800057), and to the Three Lakes Ecological Environment Protection Research and Engineering Management Center of Yuxi City for providing experimental soil.

## Data availability

Data will be made available on request.

